## Supplementary Table and Figure for "Translational reprogramming in response to accumulating stressors ensures critical threshold levels of Hsp90 for mammalian life"

Bhattacharya et al.

Supplementary Information includes one supplementary table and one supplementary figure with legend.

**Supplementary Table 1 The list of oligonucleotides used in this study.**

| **Name of oligonucleotides** | **Sequences (5´-3´)** |
|---|---|
| Human *HSP90AA1 (HSP90$\alpha$)* gRNA for CRISPR | GGTTGAGACGTTCGCCTTTC [1] |
| Human *HSP90AB1 (HSP90$\beta$)* gRNA for CRISPR | ATTGCTATTTATTCCTCGTC [1] |
| Mouse *Hsp90aa1 (HSP90$\alpha$)* genotyping forward primer | GCTGTTATGGAAGCCTCAGC |
| Mouse *Hsp90aa1 (HSP90$\alpha$)* genotyping forward primer | CACTCCAACCTCCGCAAACTC |
| Mouse *Hsp90aa1 (HSP90$\alpha$)* genotyping reverse primer | AGGGTTGTTCTCGGGACTTT |
| Mouse *Hsp90ab1 (HSP90$\beta$)* genotyping forward primer | CCGGCCTATGCAAGTCTGCACAGTGA |
| Mouse *Hsp90ab1 (HSP90$\beta$)* genotyping reverse primer | GTCCATGATGAACACACGGCGGACAT |
| Mouse *Hsp90ab1 (HSP90$\beta$)* genotyping reverse primer | CATGTTCAGTGCATAGCTCACTGACTCTG |
| sh-*HSP90AA1* (S2) target sequence | TACTTGGAGGAACGAAGAATA [2] |
| sh-*HSP90AA1* (S3) target sequence | GTTATCCTACACCTGAAAGAA [2] |
| sh-*HSP90AB1* (S2) target sequence | CGCATGGAAGAAGTCGATTAG [2] |
| sh-*HSP90AB1* (S3) target sequence | CTTGTGTTGAAGGCAGTAAAC [2] |
| sh-Control target sequence | CCTAAGGTTAAGTCGCCCTCG [2] |
| Human *GAPDH* forward primer for qRT-PCR | GCACAACAGGAAGAGAGAGACC |
| Human *GAPDH* reverse primer for qRT-PCR | AGGGGAGATTCAGTGTGGTG |
| Human *HSP90AA1* forward primer for qRT-PCR | CATAACGATGATGAGCAGTACGC |
| Human *HSP90AA1* reverse primer for qRT-PCR | GACCCATAGGTTCACCTGTGT |
| Human *HSP90AB1* forward primer for qRT-PCR | GGGTATCGGAAAGCAAGCCT |
| Human *HSP90AB1* reverse primer for qRT-PCR | ATGAGGGACATGAGTTGGGC |
| Human *CDKN1A* forward primer for qRT-PCR | AGGTGGACCTGGAGACTCTCAG |
| Human *CDKN1A* reverse primer for qRT-PCR | TCCTCTTGGAGAAGATCAGCCG |
| Human *CDKN2A* forward primer for qRT-PCR | CTCGTGCTGATGCTACTGAGGA |
| Human *CDKN2A* reverse primer for qRT-PCR | GGTCGGCGCAGTTGGGCTCC |
| Human *CDKN2B* forward primer for qRT-PCR | ATAAGGAAGCGACCTGCAACCG |
| Human *CDKN2B* reverse primer for qRT-PCR | TTCTTGGGCGTCTGCTCCACAG |
| Mouse *Gapdh* forward primer for qRT-PCR | AGGTCGGTGTGAACGGATTTG |
| Mouse *Gapdh* reverse primer for qRT-PCR | GGGGTCGTTGATGGCAACA |
| Mouse *Hsp90aa1* forward primer for qRT-PCR | GACGCTCTGGATAAAATCCGTT |
| Mouse *Hsp90aa1* reverse primer for qRT-PCR | TGGGAATGAGATTGATGTGCAG |
| Mouse *Hsp90ab1* forward primer for qRT-PCR | AAACAAGGAGATTTTCCTCCGC |
| Mouse *Hsp90ab1* reverse primer for qRT-PCR | CGTCAGGCTCTCATATCGAAT |
| Mouse *Cdkn1a* forward primer for qRT-PCR | GTCAGGCTGGTCTGCCTCCG |
| Mouse *Cdkn1a* reverse primer for qRT-PCR | CGGTCCCGTGGACAGTGAGCAG |
| Mouse *Cdkn2a* forward primer for qRT-PCR | CCCAACGCCCCGAACT |
| Mouse *Cdkn2a* reverse primer for qRT-PCR | GCAGAAGAGCTGCTACGTGAA |

| | |
|---|---|
| Mouse *Cdkn2b* forward primer for qRT-PCR | AGCAGTGTCCAGGGATGAGGAA |
| Mouse *Cdkn2b* reverse primer for qRT-PCR | TTCTTGGGCGTCTGCTCCACAG |
| Mouse *Il1b* forward primer for qRT-PCR | TGGACCTTCCAGGATGAGGACA |
| Mouse *Il1b* reverse primer for qRT-PCR | GTTCATCTCGGAGCCTGTAGTG |
| Mouse *MERVL* forward primer for qRT-PCR | TTTCTCAAGGCCCACCAATAGT [3] |
| Mouse *MERVL* reverse primer for qRT-PCR | GACACCTTTTTTAACTATGCGAGC [3] |
| Mouse *IAPEz* forward primer for qRT-PCR | GCACCCTCAAAGCCTATCTTA [3] |
| Mouse *IAPEz* reverse primer for qRT-PCR | TCCCTTGGTCAGTCTGGATTT [3] |
| Mouse *Ahsa1* forward primer for qRT-PCR | CGCCACCAACGTCAACAAC |
| Mouse *Ahsa1* reverse primer for qRT-PCR | GGCCAGGAACAGGGTTTTCA |
| Mouse *Dnajb1* forward primer for qRT-PCR | TTCGACCGCTATGGAGAGGAAG |
| Mouse *Dnajb1* reverse primer for qRT-PCR | CCGAAGAACTCAGCAAACATGGC |
| Mouse *Hspa1* forward primer for qRT-PCR | ACAAGTCGGAGAACGTGCAGGA |
| Mouse *Hspa1* reverse primer for qRT-PCR | GTTGTCCGAGTAGGTGGTGAAG |
| Mouse *Hspa8* forward primer for qRT-PCR | CCGATGAAGCTGTTGCCTATGG |
| Mouse *Hspa8* reverse primer for qRT-PCR | CCAAGGGAAAGAGGAGTGACATC |
| Mouse *Ptges3* forward primer for qRT-PCR | GGAAAGACTGGGAGGATGACTC |
| Mouse *Ptges3* reverse primer for qRT-PCR | TCATCTGCTCCATCTACTTCTGG |
| Mouse *Stip1* forward primer for qRT-PCR | TGAGTGCTGGGAACATTGATG |
| Mouse *Stip1* reverse primer for qRT-PCR | AGTCTCCTTTCTTGGCGTAGG |
| Mouse *Actb* forward primer for qRT-PCR | GGCTGTATTCCCCTCCATCG |
| Mouse *Actb* reverse primer for qRT-PCR | CCAGTTGGTAACAATGCCATGT |

**Supplementary Fig. 1 Schematic representation of the FACS gating and analysis strategies. a,** FACS gating and analysis strategy for the propidium iodide (PI) positive population of dead cells; related to Fig. 5g, and Extended Data Figs. 7b-d,f, and 9b,d. **b,** FACS gating and analysis strategy of cell cycle experiments; related to Extended Data Fig. 7a.

Supplementary Figure 1

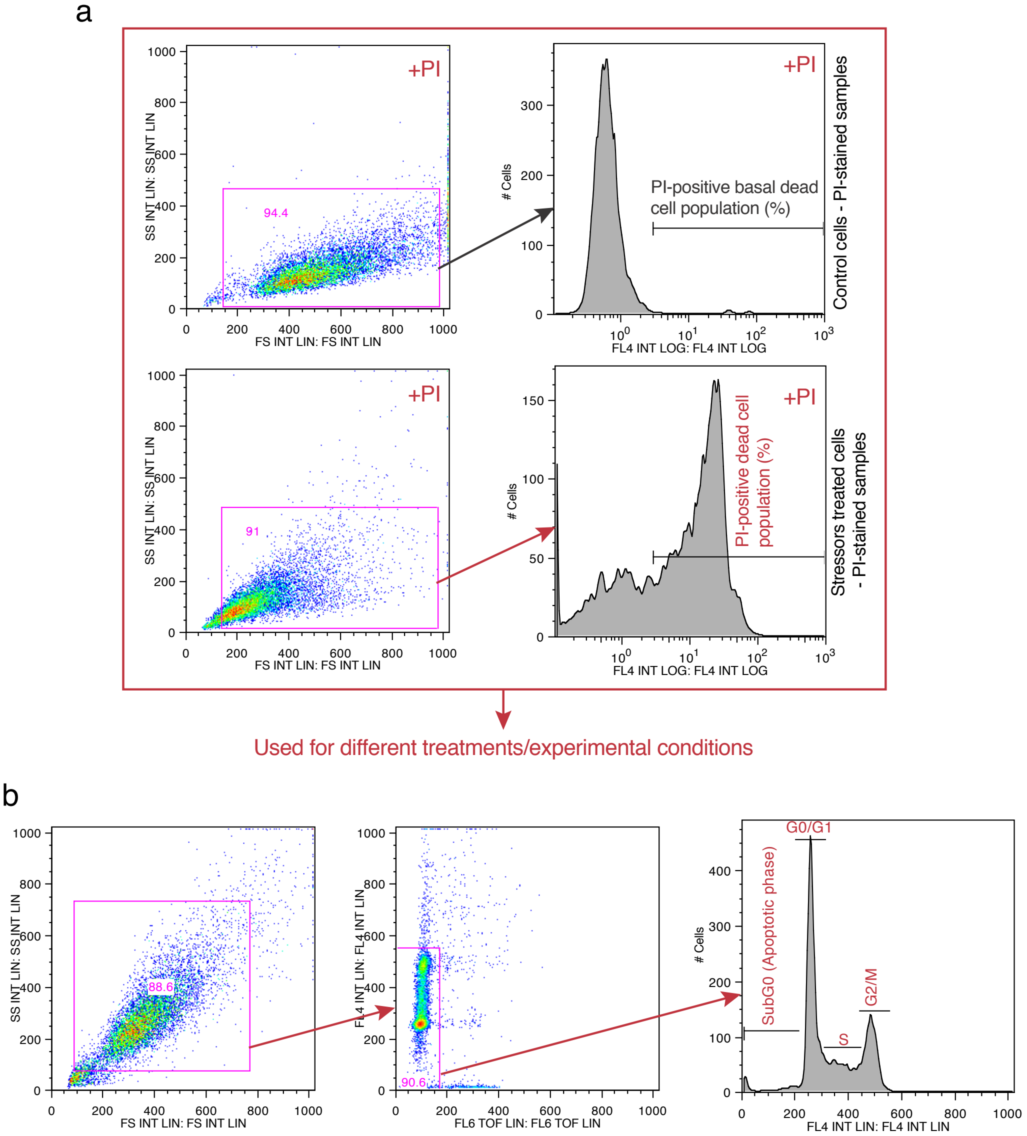
